# A comparison of ImageJ and machine learning based image analysis methods to measure cassava bacterial blight disease severity

**DOI:** 10.1101/2022.04.25.488914

**Authors:** Kiona Elliott, Jeffrey C. Berry, Hobin Kim, Rebecca S. Bart

## Abstract

**Background:** Methods to accurately quantify disease severity are fundamental to plant pathogen interaction studies. Commonly used methods include visual scoring of disease symptoms, tracking pathogen growth *in planta* over time, and various assays that detect plant defense responses. Several image-based methods for phenotyping of plant disease symptoms have also been developed. Each of these methods has different advantages and limitations which should be carefully considered when choosing an approach and interpreting the results.

**Results:** In this paper, we developed two image analysis methods and tested their ability to quantify different aspects of disease lesions in the cassava-*Xanthomonas* pathosystem. The first method uses ImageJ, an open-source platform widely used in the biological sciences. The second method is a few-shot support vector machine learning tool that uses a classifier file trained with five representative infected leaf images for lesion recognition. Cassava leaves were syringe infiltrated with wildtype *Xanthomonas*, a *Xanthomonas* mutant with decreased virulence, and mock treatments. Digital images of infected leaves were captured overtime using a Raspberry Pi camera. The image analysis methods were analyzed and compared for the ability to segment the lesion from the background and accurately capture and measure differences between the treatment types.

**Conclusions:** Both image analysis methods presented in this paper allow for accurate segmentation of disease lesions from the non-infected plant. Specifically, at 4-, 6-, and 9-days post inoculation (DPI), both methods provided quantitative differences in disease symptoms between different treatment types. Thus, either method could be applied to extract information about disease severity. Strengths and weaknesses of each approach are discussed.

## Background

Annually 20-40% of crops are lost due to plant pests and disease (FAO 2021 [1]). Causal agents of plant disease such as bacteria, viruses, oomycetes, and fungi employ various strategies to promote pathogenesis and elicit disease susceptibility in host plants. Disease susceptibility is commonly measured by the amount of *in planta* pathogen growth, reduction in crop yield/biomass, or by scaled scoring systems that use visible disease symptoms to measure severity (Strange 2003[2], Liu 2015 [3], Guant 1995 [4], Moore 1943 [5]). Each of these methods have advantages and limitations and no single method can capture the full complexity of plant disease. For instance, it is common to introduce a small number of bacteria into a plant leaf and then quantify pathogen growth overtime (Agrios 5^th^ edition 2004 [6]). This method highly quantitative and can reveal subtle differences in virulence between related pathogen strains or mutants (Bart 2012 [7], Cohn & Bart 2014 [8], Diaz, 2018 [9]). However, this assay probes only one part of the disease cycle and provides limited insight into pathogen spread, plant symptoms or defense responses. Another common method is to visually score disease symptoms on a numerical scale (Jorge & Verdier 2002 [10]). This method can be used in lab to field level experiments, is cost effective, and does not require special techniques or tools. However, accurate identification of pathogen incited symptoms can be difficult, especially in the case of multiple biotic and/or abiotic stresses. Further, disease scores may vary among different scorers and often are not sensitive enough to capture subtle changes in disease severity (Poland and Nelson 2011 [11], Strange and Scott 2005 [12]).

In recent years, there has been an increase in the use of image-based methods to analyze and measure plant health (Gehan 2017 [13], Laflamme 2016 [14], Lobet 2017 [15]). Images can be captured through many different platforms including cell phones, imaging chambers, high-throughput phenotyping facilities, drones, and satellites (Li 2014 [16], Zhang and Zhang 2018 [17]) and many analysis platforms have also been developed, for example, ImageJ (Ferreira and Rasband 2021 [18]). Image-based phenotyping tools have been successfully developed to study a broad range of plant diseases including citrus canker (Bock 2008 [19]), grapevine powdery mildew (Bierman 2019 [20]), and cereal rust disease (Gallego-Sanchez 2020 [21]). At least in some cases, image-based phenotyping can overcome some of the limitations associated with the more traditional methods described above (Mutka and Bart 2015 [22]). For example, a study investigating *Zymoseptoria trictici* infected wheat leaves found that an ImageJ analysis method provided more reliable and reproducible measures of wheat blotch disease compared to a traditional visual scoring system (Stewart 2014 [23], Stewart 2016 [24]). However, manual image analysis based on user selection of disease lesions can also be time consuming. Some image analysis methods have incorporated machine learning techniques for improved trait identification, classification, and faster analysis of plant disease symptoms (Singh 2016 [25], Tsaftaris 2016 [26]). While machine learning has enhanced the ability to process imaging data, accurate trait classification or quantification often relies on large datasets that can be expensive to acquire. Therefore, more cost effective, few-shot image analysis tools that allow for efficient segmentation and quantification of disease symptoms are needed.

In this study, we apply image-based phenotyping to cassava (*Manihot esculenta* Crantz), a starchy storage root crop (Morgan 2016 [27]). Cassava is a hardy crop predominantly grown by smallholder farmers in South America, East Asia, and Sub-Saharan Africa (Bart and Taylor 2017 [28], Hillock 2002 [29], El-Sharkawy 2003 [30]). Cassava production is threatened by the disease cassava bacterial blight (CBB). CBB can result in complete crop loss and is present in all cassava growing regions (Howler 2013 [31], Fanuo 2017 [32], Zárate-Chaves 2021 [33]). The causal agent of CBB is *Xanthomonas axonopodis* pv. *manihotis* also referred to as *Xanthomonas phaseoli* pv. *manihoti*s (*Xam* or *Xpm)* (Constantin 2016 [34]). *Xam* infects cassava by entering through open stomata or wounds in the leaf, colonizes the surface of mesophyll cells, and spreads systemically in the plant. The first visible indicators of CBB disease are dark “water-soaked” lesions that appear on the leaf. Water-soaked lesions or spots are a common, early disease symptom of various bacterial diseases. (Aung 2018 [35]). Other CBB disease symptoms include leaf wilt, defoliation, stem browning, and eventual plant death. Like other plant pathogens, *Xam* has a repertoire of effectors that can alter the structure or function of a host cell, create a more ideal environment for pathogen colonization, and overcome plant defense mechanisms (Boch 2010 [36], Hogenhout 2009 [37]). In the *Xanthomonas* and *Ralstonia* bacterial genera, this repertoire includes specialized transcription activator-like (TAL) effectors (Bodnar 2013 [38], Van Schie and Takken 2014 [39], Koseoglou 2021 [40]). TAL effectors are secreted into the plant cell and induce expression of plant susceptibility (S) genes that enhance disease. In many pathosystems, TAL effectors target *SWEET* (*Sugars Will Eventually be Exported Transporters*) genes and preventing this interaction reduces disease symptoms (Li 2012 [41], Phillips 2017 [42], Cox 2017 [43]).The *Xam* strain used in this study, *Xam668,* carries the effector, TAL20, which induces ectopic expression of *MeSWEET10a* (Cohn and Bart 2014 [8])*. Xam668* mutants with loss of TAL20 (*Xam668ΔTAL20*) exhibit visibly reduced water-soaked lesions compared to wild-type *Xam*. Here, we develop and compare ImageJ and machine learning based image analysis tools that allow for segmentation and quantification of CBB induced water-soaked lesions.

## Results

### Xam induction of water-soaked lesions in cassava

In cassava, water-soaked lesions appear as dark angular spots at the site of infection and spread as the bacteria proliferate (**Figure 1A**). To capture the progression of water-soaking in cassava, leaves were syringe-infiltrated with *Xam668*, *Xam668ΔTAL20,* or mock treatments. At 0-, 4-, 6-, and 9-days post inoculation (DPI) infected leaves were detached from the plant and imaged. Images were taken with a Raspberry Pi camera in an enclosed box to increase uniformity of imaging. An X-Rite ColorChecker Passport was included in every image for post-acquisition gray balance color correction (Berry 2018 [44]). At 4DPI, water-soaked spots began to appear in both *Xam668* (Xam WT) and *Xam668ΔTAL20* (XamΔ*TAL20*) infiltration sites (**Figure 1B**). Water-soaked lesions spread and increased in visibility at 6 and 9 DPI. However, as previously reported [33] water-soaking appeared reduced in *Xam668ΔTAL20* infection sites as compared to wildtype *Xam668* sites. Additionally, *Xam668ΔTAL20* infection sites appeared lighter in color compared to the darker lesions that develop at wildtype *Xam668* sites. Water-soaked lesions were not observed for any time point in mock infiltrated spots.

**Figure 1:**
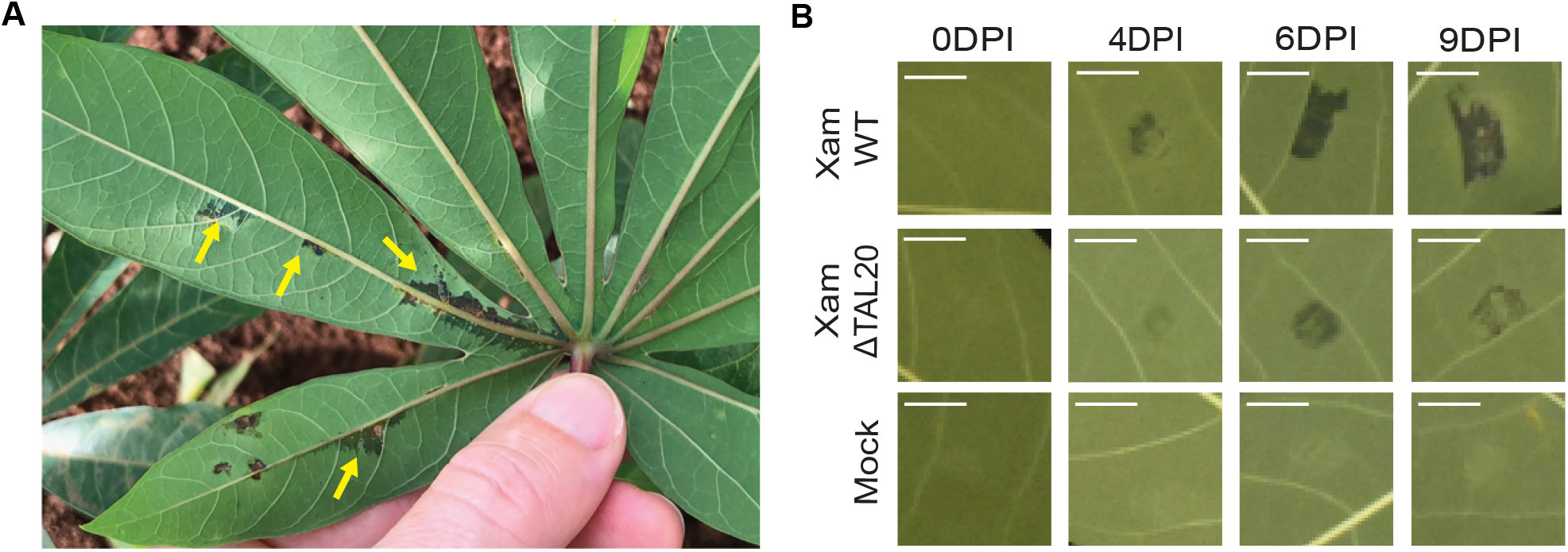
*Xanthomonas* causes complex water-soaking symptoms in cassava. **A)** Image of cassava leaf in the field exhibiting water-soaking symptoms characteristic of cassava bacterial blight. Yellow arrows indicate different water-soaked lesions. **B)** Water-soaked symptoms of cassava infiltrated with *Xam668 (*Xam WT) and a *Xam668* deletion mutant lacking the TAL20 effector (Xam*ΔTAL*20) at 0, 4, 6, and 9DPI. Mock inoculations of 10mM MgCl_2_ at each timepoint were included as controls. Scale bar = 0.5cm.

### Image J based quantification of water-soaked symptoms

ImageJ is regularly used for image analysis in biological studies (Ferreira and Rasband 2021 [18]). Here, we applied ImageJ based analysis to extract, quantify, and examine water-soaked lesion traits. Water-soaked lesions induced by *Xam668* and *Xam668ΔTAL20* were segmented using a manual overlay segmentation strategy (**Figure 2A**). For segmentation, color corrected images were uploaded and duplicated in ImageJ and the *Xam668* and *Xam668ΔTAL20* lesions were outlined using the pencil tool. Outlined images were converted from RGB to the LAB color space and the “A Channel” was obtained for better separation of the outlined lesions from the leaf background. The A channel images were thresholded and converted to a binary mask. The binary masks and analyze particle tool in ImageJ were used to define the *Xam668* and *Xam668ΔTAL20* infected sites and an overlay was created for each image. The overlays were applied to the RGB image and measurements for 27 traits were calculated. Mock sites were measured using the rectangle selection tool in the RGB image to capture information about “non-water-soaked” leaf background. ImageJ processing took approximately 6 minutes and 30 seconds per image. A movie example of the image J based analysis method was generated as a tutorial **(Additional File 1).**

**Figure 2:**
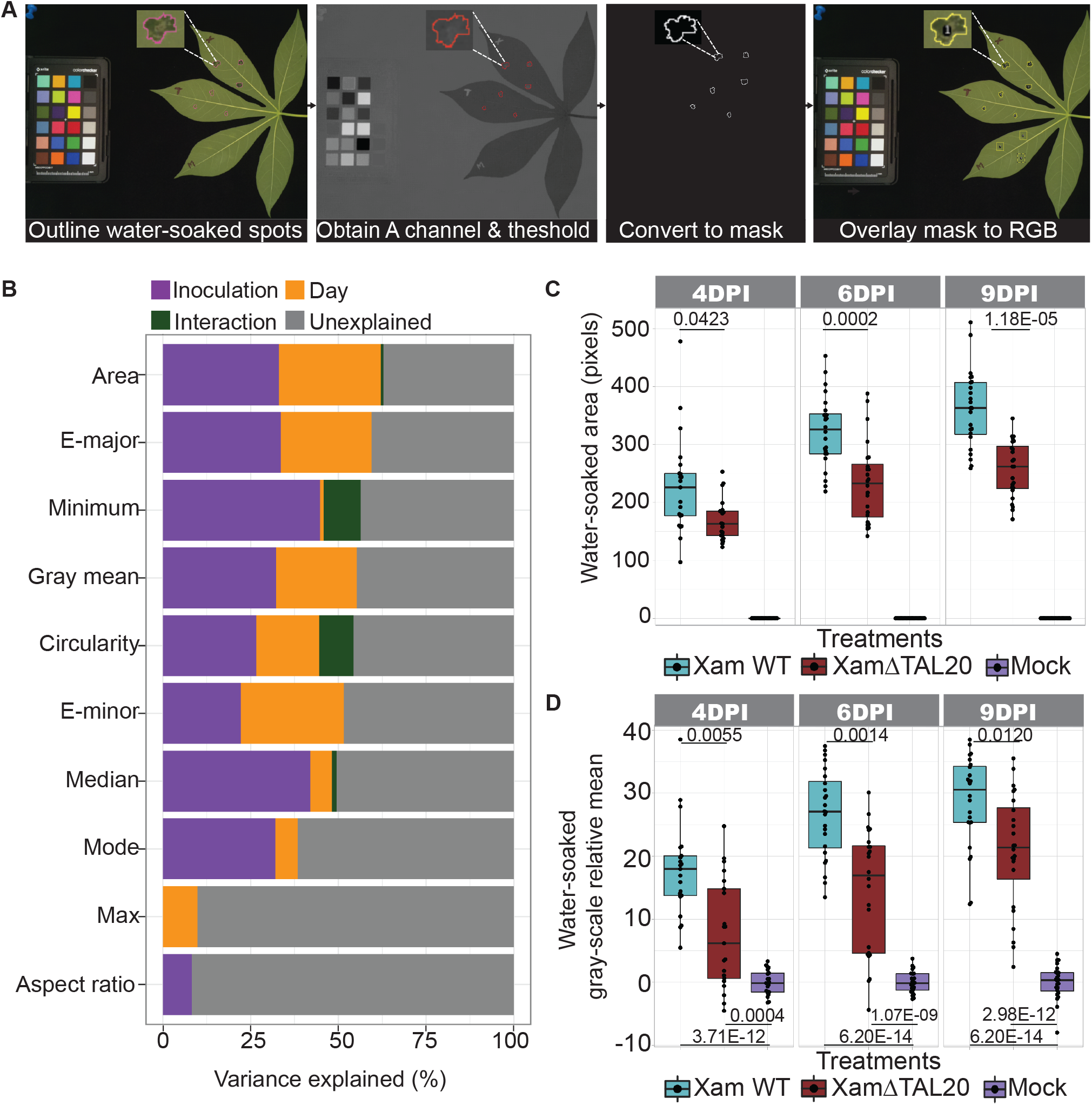
Manual ImageJ analysis of CBB water-soaking symptoms. **A)** Images of cassava leaves infiltrated with Xam WT, Xam*ΔTAL20,* and mock treatments were segmented and analyzed using an ImageJ overlay segmentation method. Overlay segmentation analysis depicted by step using a CBB infected cassava leaf image. Images were taken at 0, 4, 6 and 9 DPI. Leaf lobes were labeled by treatment type: X=Xam WT, T=Xam*ΔTAL20*, and M=Mock. White lines point to selected regions of a representative water-soaked lesion at each step of the ImageJ overlay segmentation process. **B)** The variance explained by inoculation type (Xam WT or Xam*ΔTAL20*), DPI (4-, 6- and 9-), or the interaction between inoculation type and DPI for ten ImageJ generated measurements. Variances were determined by ANOVA analysis. **C)** Total water-soaked area (pixels, y-axis) for sites infiltrated with each treatment (x-axis). Calculated *p*-values (Kolmogorov-Smirnov test) shown above the line in each plot. **D)** Negative gray-scale mean (y-axis) of water-soaked lesions for Xam WT and Xam*ΔTAL20* relative to mock inoculated spots (x-axis) within the same leaf. Calculated *p*-values (Kolmogorov-Smirnov test) shown above the line in each plot. In ImageJ, the gray-scale mean was measured by averaging the mean of each gray-scale value in the RGB channels.

Ten traits were selected and further analyzed using an ANOVA analysis to determine the variance explained (VE) by three terms of interest: (1) inoculation type, (2) DPI and (3) the interaction between inoculation type and DPI (**Figure 2B**). Inoculation type and DPI were selected as defining factors because we expected that water-soaking severity is dependent on these terms. Area had the highest amount of VE, with over 60% VE. We selected gray-scale mean as another trait of interest because of the color difference we observed between *Xam668* and *Xam668ΔTAL20* water-soaked lesions. Gray-scale mean, accounted for over 50% VE. Water-soaked area (**Figure 2C)** and gray-scale mean (**Figure 2D)** were further analyzed as measures of CBB disease severity. The *Xam668* sites had significantly more water-soaked area compared to *Xam668ΔTAL20* at each timepoint. We found there was noise in the gray-scale mean data due to lack of standardization across individual images despite gray balance color correction. To account for this, a linear model was applied to determine the grand mean of all gray values in each image and the *Xam668* and *Xam668ΔTAL20* gray values were centered to mock. In each timepoint, *Xam668* treatment resulted in lesions that had a significantly larger gray-scale mean compared to *Xam668ΔTAL20* treatment. A greater difference in gray-scale mean was observed between *Xam668* and mock treated spots compared to *Xam668ΔTAL20* and mock spots. These results indicate that ImageJ based segmentation allowed for separation of treatment types and for the quantitative analysis of water-soaked lesions over time.

### Machine learning based quantification of water-soaked symptoms

While ImageJ provided sufficient segmentation of water-soaked lesions, developing an overlay mask for every individual image is time intensive. Therefore, we sought to develop a machine learning tool that would provide faster segmentation and quantification of diseased leaves. A custom workflow for machine learning disease lesion analysis was developed using the source file from PhenotyperCV, a C++11 library designed for image-based phenotyping (Berry 2018 [44]). The machine learning workflow was run using the Mac terminal. Command syntax specific for each step of the machine learning tool was developed **(Additional File 2)**. Five representative images of CBB infected leaves from different DPI were selected and combined into one graphic as a training image for the machine learning tool (**Figure 3A**). A binary mask was generated from the combined leaf graphic using ImageJ. The mask was used to generate a support vector machine (SVM) learning classifier (YAML) file. The classifier file was used to process the images and eliminated the need to manually outline each lesion or make individual masks (**Figure 3B**). During processing, images were color corrected and manually thresholded using a scale bar built into the program to reduce background noise and enhance segmentation of lesion pixels. Next, infiltrated spots were manually labelled and color-coded by treatment type. Output images were generated and included color corrected, pseudo-color map, and feature prediction images for every image analyzed (**Figure 3C**). Machine learning processing took approximately 2 minutes and 30 seconds per image. Processing speed increased when all images were analyzed using an iteration (for loop) command in terminal allowing the machine learning tool to be executed on several images in succession. A movie example of the machine learning based analysis method was generated as a tutorial **(Additional File 3).** Additionally, two space separated text (TXT) files were produced with shape and color related measurements of each lesion. A list of the reported measurements is included **(Additional File 4)**. Shape data generated by the machine learning tool includes area, hull area, height, width, etc. The color data generated by machine learning is a lightness histogram of 0-255 for each lesion which was used to calculate lesion gray-scale mean.

**Figure 3:**
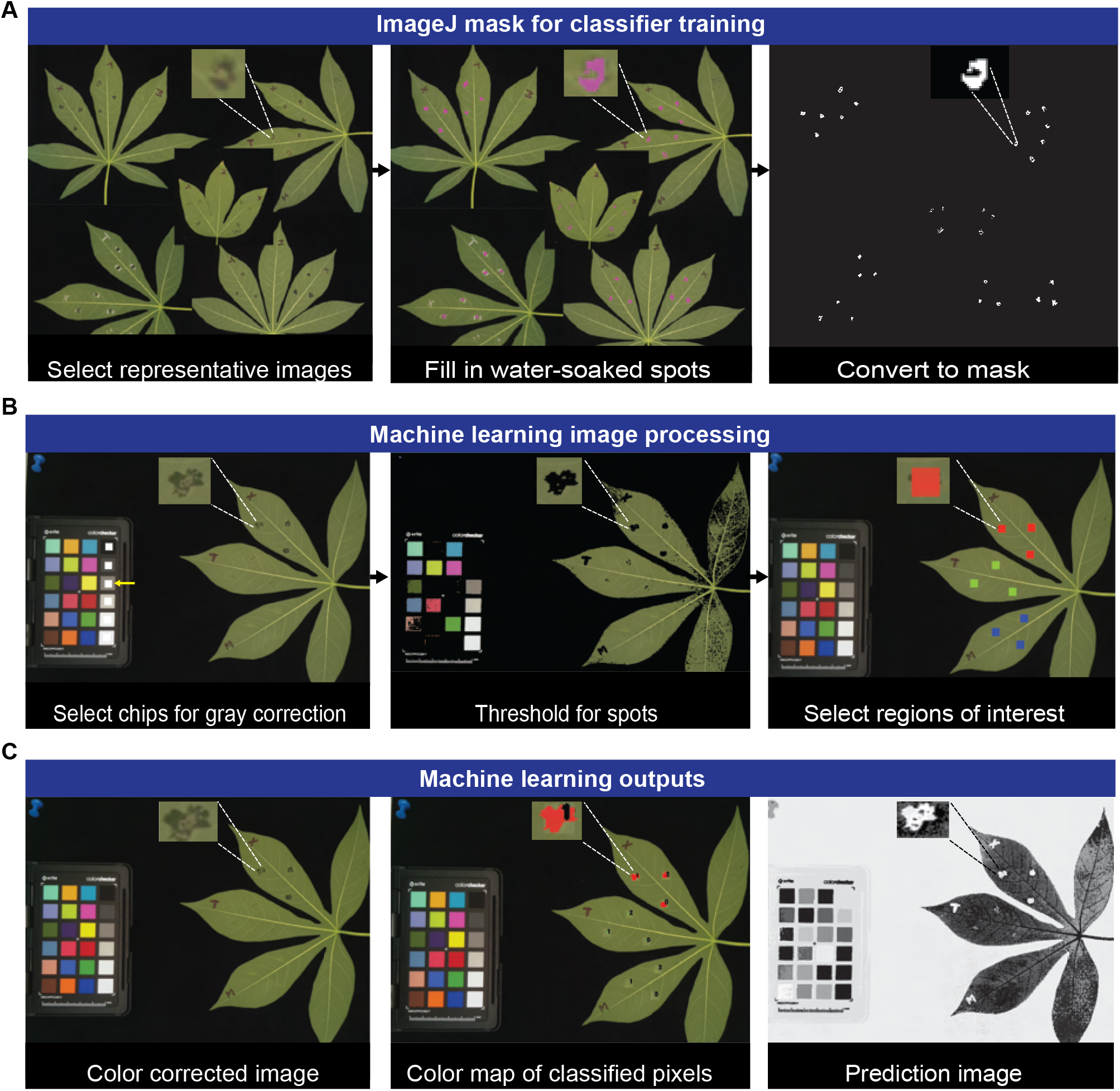
Overview of the Support Vector Machine learning segmentation and analysis method. **A)** Images of cassava leaves infiltrated with Xam WT, Xam*ΔTAL20,* and mock treatments were segmented and analyzed using a support vector machine learning tool. Images depict steps used to generate a classifier training mask for the machine learning tool. A mask was made by combining representative CBB infected images into one graphic and generating a binary mask in ImageJ. White lines showcase a representative water-soaked lesion within the combined leaf graphic and indicate changes at each step. The mask was used to generate a classifier (YAML) file with PhenotyperCV. **B)** Images depict steps of machine learning processing using a CBB infected cassava leaf image. Images were uploaded into the machine learning tool and processed by gray balance color correction, thresholding, and the inoculated regions of interest were selected and labeled using a color code: Red=Xam WT, Green= Xam*ΔTAL20* and Blue= Mock. White lines showcase a representative water-soaked lesion within the image and indicate changes at each step. **C)** Images exhibit outputs from the machine learning image processing and include the color corrected image (left), a pseudo-colored map of the pixels classified as water-soaked (middle), and a feature prediction image (right). White lines showcase a representative water-soaked lesion within the image and indicate differences in each output image. Text separated files with shapes and color data for each inoculation spot were also generated.

Twelve machine learning derived traits were selected and the ANOVA analysis was used to measure VE by each trait (**Figure 4A**). Area measured by the machine learning tool had over 75% VE by the defining factors. As was determined during ImageJ analysis, area also accounts for the highest amount of VE in the machine learning analysis. The gray-scale mean had over 60% VE by the defining factors. Consistent with the ImageJ analysis, the machine learning approach revealed that *Xam668* caused a larger water-soaked area (**Figure 4B)** and relative gray-scale mean (**Figure 4C)** compared to *Xam668ΔTAL20* infiltrated spots. These data suggest that the machine learning tool adequately distinguished between treatment types and provided quantitative measures of water-soaked lesions using the classifier file created from one training mask.

**Figure 4:**
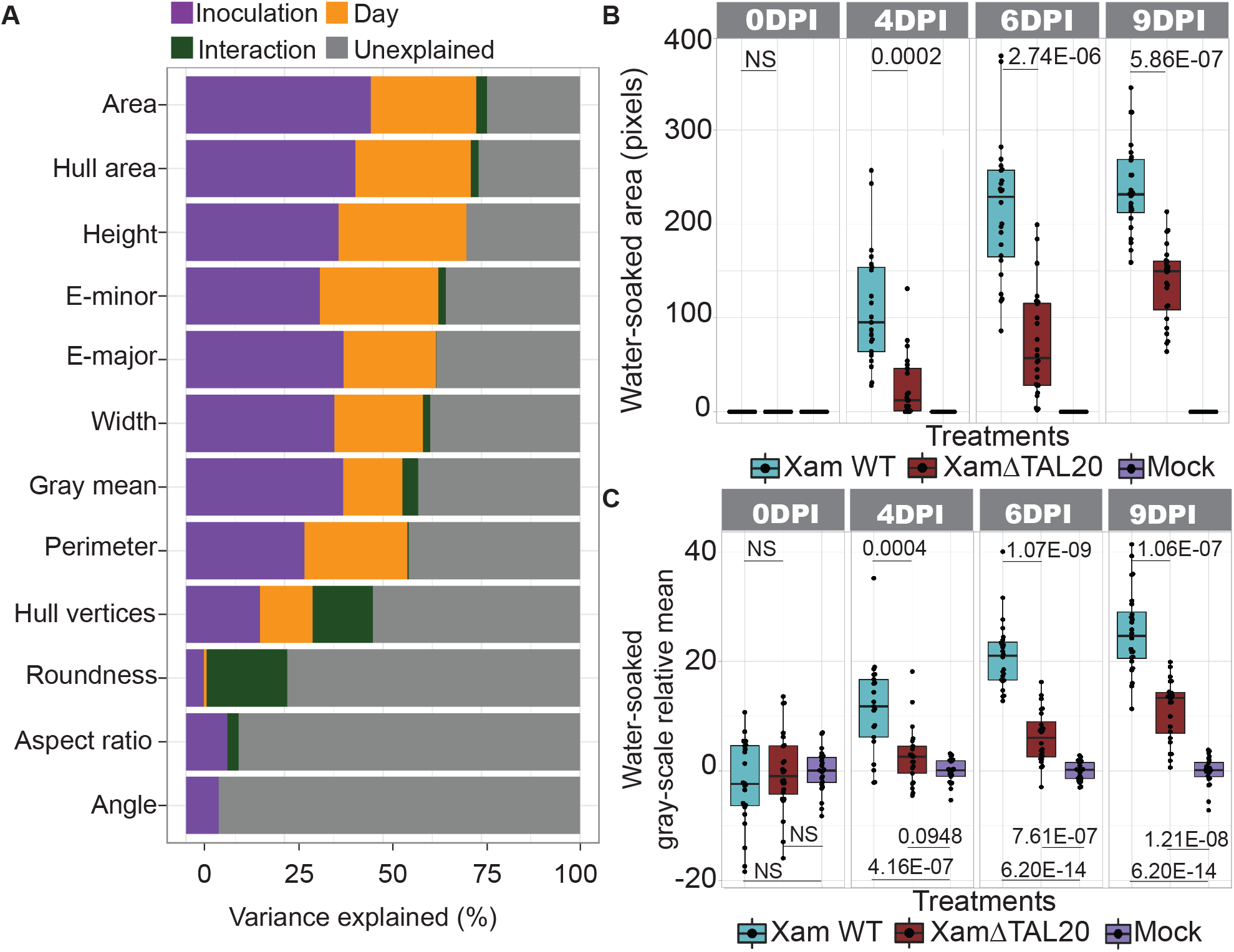
Support Vector Machine learning analysis of CBB water-soaked symptoms. **A)** The variance explained by inoculation type (Xam WT or Xam*ΔTAL20*), DPI (4-, 6-and 9-), or the interaction between inoculation type and DPI for twelve machine learning generated measurements. Variances were determined by an ANOVA. **B)** Total water-soaked area (pixels, y-axis) for sites infiltrated with each treatment (x-axis). Calculated *p*-values (Kolmogorov-Smirnov test) shown above the line in each plot. **C)** Negative gray-scale mean (y-axis) of water-soaked lesions for Xam WT and Xam*ΔTAL20* relative to mock inoculated spots (x-axis) within the same leaf. Calculated *p*-values (Kolmogorov-Smirnov test) shown above the line in each plot. In the machine learning analysis, the gray-scale mean was generated using the average mean of the “L” channel from the LAB color space.

### Comparison of the ImageJ and Machine learning based lesion analysis methods

The ImageJ and machine learning based methods both successfully distinguished *Xam668* and *Xam668ΔTAL20* and yet the results were not equivalent. To further compare and contrast these methods, representative *Xam668* and *Xam668ΔTAL20* lesions from 4-, 6-, and 9-DPI were selected and visually inspected (**Figure 5A**). We observed that machine learning was able to distinguish between water-soaked and “non-water-soaked” pixels within the lesion spot whereas in ImageJ, a boundary was put around the whole spot and could include a mix of both pixel types. This suggests that the machine learning tool is more selective in classification of water-soaked versus non-water-soaked pixels and would explain the trend of overall smaller area measurements generated by machine learning compared to ImageJ. In ImageJ, the lesion boundary is user-selected. However, to completely separate water-soaked from non-water-soaked pixels in lesions where there is a mix, smaller independent boundaries would be required. Having multiple boundaries for one lesion is not ideal as it would impact measures such as gray-scale mean and increase image processing time. The two image analysis methods were statistically compared by pairing the mock, *Xam668* and *Xam668ΔTAL20* area data and performing F-statistic variance tests on each respective treatment type (**Figure 5B**). At each timepoint, there was no significant difference in the variance observed between ImageJ and machine learning data suggesting the two methods have equal variation within each treatment type.

**Figure 5:**
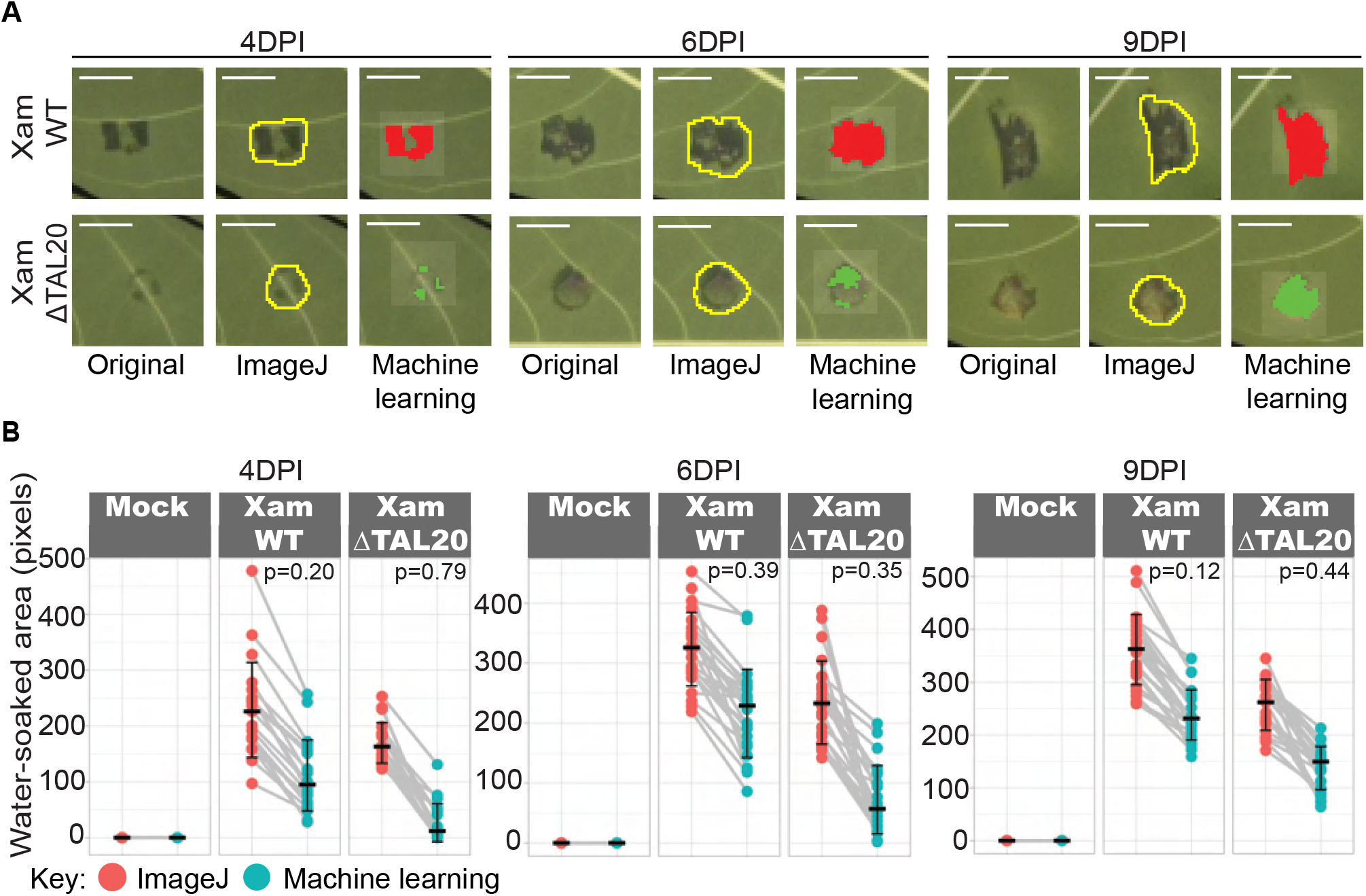
Comparison of the ImageJ and machine learning analyses of CBB infected leaves. **A)** Representative images from each timepoint (4-, 6-, and 9-DPI) of a Xam WT (top row) and Xam*ΔTAL20* (bottom row) water-soaked spots were selected, visually inspected, and compared. The original images show the water-soaked spots from the color corrected images without segmentation from the background. The “ImageJ” images show water-soaked spots manually segmented from background and overlaid onto the RGB image. The machine learning images shows water-soaked spots segmented from background and pseudo-colored. Scale bar = 0.5cm **B)** Water-soaked area data generated by ImageJ or machine learning were paired by inoculation location and plotted for 4 DPI (left plot), 6 DPI (middle plot), and 9 DPI (right plot). Calculated *p-*values (F-Variance test) shown in the upper corner of plot. Red=ImageJ Blue=machine learning.

## Discussion

To quantify CBB, we developed and compared ImageJ and machine learning image analysis methods for accurate segmentation and quantification of water-soaked lesion symptoms. We found that an ImageJ overlay segmentation method allowed for adequate separation between cassava infected with mock, *Xam668* and *Xam668ΔTAL20* treatments based area and gray-scale mean values of disease lesions. However, the ImageJ analysis was time-consuming because an individual mask had to be made for every image analyzed. Other ImageJ analysis methods tested with this data set such as non-segmentation and color-threshold based segmentation of water-soaked lesions failed to accurately capture the water-soaking phenotype.

Machine learning has previously been applied to detect and measure several cassava diseases including bacterial blight, brown streak and mosaic disease (Sangbamrung 2020 [45], Ramcharan 2019 [46]). However, these tools rely on hundreds to thousands of images for classifier training. Any machine learning tool is heavily reliant on its classifier file for adequate segmentation and measure of an object of interest. If a classifier file does not adequately capture the range of traits for an object of interest, classification of that object will fail. To determine if a classifier file would work accurately for our data set, we tested its predictive capability by spot checking analysis accuracy in a subset of images and visually inspecting classification of pixels defined as water-soaked. We initially developed classifier files based on a single representative CBB infected leaf image and found it could not reliably predict features of interest for all images. However, by combining representative images of cassava infected with three replicates each of mock, *Xam668,* and *Xam668ΔTAL20* treatments across different timepoints into one training graphic, we developed a classifier that better predicted water-soaked lesions. The accuracy of the combined leaf graphic was tested by again spot-checking a subset of color map images and inspecting classification of pixels defined as “water-soaked”. Similarly, our classifier file was developed using one genotype of cassava, TME419. In future studies, if this approach were to be applied to datasets derived from multiple genotypes or a breeding program, the classifier file would need to be updated with representative images to capture any additional variability in leaf traits.

Another important consideration for classifier file development is the machine learning algorithm used. The machine learning workflow presented here functions with either support vector machine (SVM) or Naïve Bayes learning algorithms. During testing of classifier files, we found that SVM training files predicted water-soaked lesion features in our system more accurately than Naïve Bayes. Similarly, a previous study tested three machine learning methods and reported that SVM had high performance in predicting and classifying cassava diseases (Ramcharan 2017 [47]).

Despite the limitations, we found that the few-shot machine learning based image analysis tool presented here offered a fast and accurate approach to segment water-soaked lesions. Processing for the machine learning tool took less than half the time of ImageJ based analysis for each image. The machine learning tool worked as well as the ImageJ overlay segmentation method for separating lesions by treatment type and extracting quantifiable data. Due to the time needed to validate a classifier file, we suggest that a machine learning approach for image-based lesion analysis is appropriate when there is a large number of images to be processed. If the data set is small, ImageJ could be a faster approach as the accuracy of the method does not rely on a classifier file. Moreover, manual thresholding is still required for segmentation of the lesions in each image and may be slightly variable within the data set. Thresholding performed within either the machine learning or ImageJ methods requires user decision to determine the threshold cut-off. In the case of the machine learning tool, it is important to inspect the color maps generated for each image analyzed to ensure proper classification of water-soaked lesions. In some cases, we found it necessary to re-process images in the machine learning tool and adjust the threshold for more precise capture of a lesion.

While improvement is still needed in image-based phenotyping, there are several potential uses for the machine learning and ImageJ analyses presented in this study. Image based phenotyping has become increasingly popular for examining the link between disease symptoms and genetics in plant science (Casto 2021 [48]). The tools presented here provide a new resource for experiments investigating CBB disease susceptibility. Additionally, the general framework of the machine learning workflow can be applied to other plant species and disease symptoms using classifier files representative of the disease of interest.

## Conclusions

To quantify CBB, we developed and compared ImageJ and machine learning image analysis methods for accurate segmentation and quantification of water-soaked lesion symptoms. Both the ImageJ and machine learning image analysis methods are described in detail, along with video tutorials and we hope these resources will help other researchers use these tools and/or design similar tools that can be applied to other pathosystems. We found that both methods accurately distinguished between and quantified different water-soaked lesion types in the cassava-*Xanthomonas* pathosystem. The ImageJ method is best used from smaller datasets as it relies on the user developing a mask for every image. The machine learning based tool is best used for larger datasets as it is more time efficient to develop a single classifier file to process many images. Many machine learning tools rely on thousands of training images for accurate function. However, the machine learning tool presented here is few-shot learning based and functions as well as ImageJ for disease segmentation and measurement.

## Methods

### Plant materials and growing conditions

Cassava plants from the cultivar TME204 were kept in greenhouse conditions set to 28°C; 50% humidity; 16 hrs light / 8 hrs dark and 1000 W light fixtures that supplemented natural light levels below 400 W / m^2^. Cuttings were taken from the woody stem of mature plants and propagated to 4-inch pots of Berger45 soil. 4–5-week-old propagated plants that were well established were used for infection experiments. During infection experiments, plants were kept in a post-inoculation room set to 50% humidity, ambient room temperature, 12 hrs light / 12hr dark and 32 W light fixtures.

### Bacterial inoculations

*Xanthomonas* strains were struck from glycerol stocks onto NYG agar plates containing appropriate antibiotics. The strains used for this study were *Xam668* (rifampicin 50 µg/ml) and *Xam668ΔTAL20* (suicide vector knockout (Cohn and Bart 2014 [8]) tetracycline 5 µg/ml, rifampicin 50 µg/ml). *Xanthomonas* strains were grown in a 30°C incubator for 2-3 days. Inoculum for each strain was made by transferring bacteria from plates into 10mM MgCl_2_ using inoculation loops and brought up to a concentration of OD_600_ = 0.01. Leaves from 4–5-week-old cassava plants were inoculated using a 1.0 mL needleless syringe. For each replicate assay, two cassava plants were used for inoculations and four leaves were inoculated on each plant. One bacterial strain was inoculated per leaf lobe with three injection sites. Mock inoculations of 10mM MgCl_2_ alone were included resulting in nine infiltrated sites per leaf. Four replicate rounds of inoculations were done in total.

### Imaging

Cassava leaves were detached and imaged at 0-, 4-, 6-, and 9-days post inoculation (DPI). One leaf from each cassava plant was collected and imaged for a total of two leaves per timepoint. In all, thirty-two leaves were imaged and analyzed across four replicate rounds of inoculations. Leaves were imaged from above using a Raspberry Pi Sony IMX219 camera in an enclosed box with an overhead light. To account for setting inconsistencies between images, images were color-corrected by gray balancing using a X-Rite ColorChecker Passport color card. Images were uploaded to the machine learning workflow and six gray color chips (black-white) were manually selected using a selection tool built into the program. Saturation of each chip was estimated and the brightness of each image was adjusted accordingly. The gray corrected images were then used for water-soaking analysis. Analytical standardization of the gray values post-image-processing by ImageJ and machine learning was performed separately by estimating the grand mean of all gray values within each image and centering those values to the grand mean across all images. This is achieved by creating a linear model with a single fixed effect term accounting for each image and extracting model residuals.

### ImageJ image analysis

Gray corrected images were uploaded to ImageJ version FIJI (Schindelin 2012 [49]) and duplicated. Water-soaked lesions were manually outlined on the duplicate image using the pencil tool (color: #ff00b6 and size 2). The outlined images were converted from RGB to LAB and split to obtain the A color channel. The A channel images were thresholded, converted to a mask and the mask for each spot was added to the ROI manager using the analyze particle tool. The ROI masks were applied to the original RGB gray corrected images. Mock infiltrated spots (no water-soaking, plant background data) were added to the ROI manager using an arbitrarily sized rectangle selection tool consistently set to a W=26 and H=30. Area, gray-scale mean, and eight other measurement data were obtained for each infiltrated spot using the FIJI measure tool. The measurements were saved as a comma separated value (CSV) file. The variance explained by ten image J derived traits were calculated and plotted in the software program R using a custom partial correlations script. Area and gray-scale mean data for all lesions were compared across different treatment types and timepoints using a Kolmogorov-Smirnov (KS) statistical test in R. All plots were generated in R with a dpi=300, width=8.66, and height=6.86.

### Machine learning image analysis

Five images of *Xanthomonas* inoculated cassava leaves from different timepoints were selected as representatives to make a classifier file for the machine learning image analysis tool. The images were combined into one graphic, uploaded to ImageJ, and water-soaked spots were outlined and filled in using the pencil tool (color: #ff00b6). The outlined combined leaf image was converted to a binary mask and referred to as the “labeled image”. The machine learning image analysis tool is part of PhenotyperCV, a C++11 header-only library designed for image-based plant phenotyping. The machine learning workflow and software download instructions are available on GitHub (https://github.com/jberry47/ddpsc_phenotypercv/wiki/Machine-Learning-Workflow).

All steps of the machine learning workflow were run on the Mac terminal command line. The labeled leaf mask image and original combined leaf graphic were used to create a support vector machine learning classifier or YAML file. Individual images of inoculated cassava leaves were processed in the machine learning tool by uploading the images and gray correcting. The images were thresholded using a scale bar built into the program to set a cut-off for pixels that can be classified as water-soaked. The inoculated sites were manually selected with a color-coded region of interest (ROI) selector (mouse right click-red, left click-green, and middle click-blue). The ROI selector tool size ranges from 0-20. The ROI size was consistently set to 11 for this study. The ROI selector does not restrict the size of the object identified as a water-soaked lesion. If a part of the object defined as a lesion is included in the ROI selection, then the entire object will be labelled and color-coded. For this study, we designated red as *Xam668*, green as *Xam668ΔTAL20,* and blue as mock inoculation spots. If color-code separation is not required for other studies using the machine learning tool, one click/color type can be used for all lesion selections. Outputs from the workflow include a color corrected image (also used in the ImageJ analysis), a prediction image of what could be captured as pixels of interest, and a pseudo-colored map image showing what was captured as pixels of interest. Additionally, two space separated text files were generated with measurement data about the shape and color of each lesion. The shape file includes nineteen trait measures such as area, height, circularity, etc. The color file includes is a lightness histogram of 0-255 for each lesion. The text files were uploaded into R and processed using a custom script designed to read and format the data and create a comma separated value (CSV) file. For the color file, the histogram data were used to calculate lesion gray-scale mean. The variance explained by twelve machine learning derived traits were calculated and plotted in R using a custom partial correlations script. Area and gray-scale mean data for all lesions were compared across different treatment types using a Kolmogorov-Smirnov (KS) statistical test in R. All plots were generated in R with a dpi=300, width=8.66, and height=6.86.

## List of abbreviations

CBB: Cassava Bacterial Blight
Xam: *Xanthomonas axonopodis* pv. *manihotis*
Xpm: *Xanthomonas phaseoli* pv. *manihotis*
PV: Pathovar
DPI: Days post inoculation
TAL: Transcription activator like
VE: Variance explained
SVM: Support vector machine learning
WT: Wildtype
SWEET: Sugars will eventually be exported transporters
ROI: Region of Interest
KS: Kilmogrov-Smirnov
CSV: Comma separated plain text file

## Declarations

### Ethics approval and consent to participate

Not applicable

### Consent for publication

Not applicable

## Availability of data and materials

The datasets and custom R scripts generated and/or analyzed in this study are available in the figshare repository, https://figshare.com/s/0148e5e4fc7f220ac4c3

## Authors’ contributions

KE and RSB designed the study. KE completed bacterial inoculations and collected images at each timepoint. HK and KE completed ImageJ analysis and interpreted the results. JB set up the Raspberry Pi camera system and developed the machine learning workflow in PhenotyperCV. JB and KE tested the machine learning tool performance. KE processed all images analyzed through the machine learning tool and interpreted results. JB developed the gray correction and image effects color correction methods for image analysis. JB provided expertise on statistical analyses and developed initial R scripts used to run statistical tests. KE wrote the original manuscript draft, completed statistical analysis, and generated figures. KE, JB, and RSB assisted in manuscript and figure review and editing. RSB provided supervision over the project. All authors reviewed and approved the manuscript.

## Acknowledgements

We acknowledge the Bart Lab members who provided insightful discussion and feedback on this project, especially Dr. Kira Veley, Dr. Qi Wang, Dr. Ben Mansfeld, and Taylor Harris.

## Authors’ information

**Kiona Elliott:** PhD candidate at Washington University in St. Louis and the Donald Danforth Plant Science Center.

**Jeffrey C. Berry:** Sr. Data Scientist at the Donald Danforth Plant Science Center

**Hobin Kim:** Donald Danforth Plant Science Center high summer school intern from the Army and Navy Academy.

**Rebecca S. Bart:** Associate member and principal investigator at the Donald Danforth Plant Science Center.

## Supplementary

**Additional File 1: Movie example of ImageJ based analysis method** Available for download on figshare or online at https://youtu.be/EtEzRls4Jh4

**Additional File 2: Machine learning tool commands. A table of the command syntax, function, and description of inputs and outputs for each command.**

**and description of inputs and outputs for each command.**
| Tool Commands: | Function: | Inputs Needed: | Outputs: |
| --- | --- | --- | --- |
| ML_CREATE | Create .YAML classifier file based on of Naïve Bayes or Support Vector Machine learning algorithms | Representative RGB (input) and mask (labeled) images | .YAML file |
| ML_PRED | Create prediction image of what pixels will be classified as object of interest based on classifier file | Classifier file, input image, and type of machine learning method used to create classifier file | Prediction .PNG file |
| ML_PROC | Process all images using appropriate classifier type | Classifier file, input image, and type of machine learning method used to create classifier file, size of ROI selection box | Gray corrected, color-map and prediction .PNG file for each image and .txt files with shape and color data for each object |
| ML_STAT | Create summary statistics of classifier file accuracy rate | Classifier file, input image, and type of machine learning method used to create classifier file | Summary true positive, false positive, false negative, true negative, precision, recall, accuracy, true positive rate, false positive rate, true negative rate, and false discovery rate statistics for the classifier file |

**Additional File 3: Movie example of machine learning based analysis method** Available for download on figshare or online at https://youtu.be/Dw2VebjExZw

**Additional File 4: Machine learning measurement types. A table of measurements generated from the machine learning tool and their descriptions**

| **Measurement type:** | **Description:** |
| --- | --- |
| Area | Area of object |
| Hull area | Area of convex hull ( shape of the smallest convex set) |
| Solidity | Ratio: area / convex hull area |
| Perimeter | Perimeter around object |
| Width | Width of object |
| Height | Height of object |
| Cmx | Center of mass x: position of center of mass |
| Cmy | Center of mass y: position of center of mass |
| Hull verticies | Number of convex hull vertices |
| Ex | Ellipse center x: position of the center of the minimum bounding ellipse |
| Ey | Ellipse center y: position of the center of the minimum bounding ellipse |
| E-major | Ellipse major axis: length of the major axis of the minimum bounding ellipse |
| E-minor | Ellipse minor axis: length of the minor axis of the minimum bounding ellipse |
| E-angle | Ellipse angle: angle of rotation of the bounding ellipse major axis |
| Eccen | Ellipse eccentricity: eccentricity of the bounding ellipse |
| Circ | Circularity |
| round | Roundness |
| Ar | Aspect ratio |
| Fd | Box count fractal dimension |
| Gray scale | Lightness channel histogram from 0-255 |

